# Dynamic corticostriatal neural coordination governs competition between social and metabolic drives in rats

**DOI:** 10.64898/2026.01.19.700365

**Authors:** Rishika Tiwari, Shoval Raz, Shai Netser, Shlomo Wagner

## Abstract

Animals must continuously prioritize among competing internal needs, yet the neural mechanisms that arbitrate between these drives remain poorly understood. Here, we used a behavioral paradigm in rats to quantify choice between social interaction and food seeking under varying need states. We show that social isolation selectively shifts preference toward social engagement, even under metabolic demand, indicating a reweighting of motivational priorities. Chronic local field potential recordings across prefrontal and striatal brain regions revealed that this shift is accompanied by a reconfiguration of corticostriatal theta-band coordination. Trial-by-trial analyses demonstrated that network dynamics predict behavioral choice, linking neural coordination to decision output. Pharmacological blockade of oxytocin receptors attenuated both the network reorganization and the behavioral prioritization shift, implicating oxytocin signaling in gating this process. Together, our findings identify a network-level mechanism through which internal states dynamically compete, providing a framework for understanding how the brain resolves conflicting motivational demands.

## Introduction

Animals must continuously prioritize among competing internal needs, such as hunger, thirst, and social interaction. While the neural circuits underlying individual motivational drives have been extensively studied (Flavell et al., 2022), far less is known about how the brain arbitrates between concurrent needs to guide behavior.

Social interaction represents a fundamental biological need, and its deprivation produces robust behavioral changes across species (Lee et al., 2021). However, the neural mechanisms by which social need is integrated with other homeostatic drives, such as energy balance, remain poorly understood. In particular, it is unclear how internal states reshape network dynamics in the brain to bias behavioral choice under conditions of competing motivation.

Motivational behaviors are driven in part by the brain’s reward system, whereby increasing need enhances the rewarding value of its fulfillment (Brehm & Self, 1989; Isaac & Murugan, 2024). In rodents, both food and social interactions are rewarding and engage overlapping corticostriatal circuits, including the nucleus accumbens (Acb), orbitofrontal cortex (OFC), and medial prefrontal cortex (mPFC) (Bein & Niv, 2025; Borland, 2024; Jabarin et al., 2025; Marinescu & Labouesse, 2024).

Oxytocin is another key regulator of motivation. This hypothalamic neuropeptide modulates both social and feeding behaviors and acts broadly across reward- and social-processing circuits (Dölen et al., 2013; Gordon et al., 2011; Onaka & Takayanagi, 2019). However, whether oxytocin signaling mediates isolation-induced shifts in motivational trade-offs between social interaction and food seeking remains unclear.

Here, we used a behavioral paradigm quantifying choice between social interaction and food seeking under varying need states (Netser et al., 2020; Reppucci et al., 2020). Combining this approach with chronic recordings across prefrontal and striatal regions, we examined how social isolation alters neural coordination during motivated decision-making. We further tested the role of oxytocin signaling in modulating these dynamics. Our results identify a network-level mechanism by which internal states reconfigure corticostriatal activity to bias behavioral prioritization, providing insight into how the brain resolves competition between fundamental motivational drives.

## Materials and methods

### Animals

Adult male and female Sprague–Dawley rats (12–16 weeks; subjects) were bred in-house or obtained from Envigo (Israel) and served as subjects, while Sex-matched juveniles (4–7 weeks) served as social stimuli. Animals were housed under a reversed 12 h light/dark cycle with *ad libitum* access to food and water unless otherwise specified. All procedures were approved by the University of Haifa IACUC (#1664U, #2059U) and followed NIH guidelines.

### Surgical implantation of electrode array

Custom-made electrode arrays (EArs) were fabricated as previously described (Mohapatra et al., 2022). Animals were anesthetized by intraperitoneal administration of Ketamine and Domitor (0.09 mg/g and 0.0055 mg/g body weight of the rat, respectively). Sedation of animals was confirmed by toe pinch reflexes. The body temperature of the animals was kept constant at approximately 37°C using a custom-made temperature controller connected to a heating pad placed under the animal. Anesthetized animals were fixed in a stereotaxic apparatus (Stoelting Inst.), keeping the head flat using ear bars. The eyes were covered by Duratears ointment (Alcon couvreur N.V., Belgium) to prevent drying of the cornea. To maintain sedative state, isoflurane (0.1-0.5%) was constantly supplied to the nose (Somnoflow® low-flow electronic anaesthesia system, Kent scientific). Skull skin was removed carefully using a scalpel and scissors, while cold saline was applied to the incised area to clean it. Craniotomy was then performed, and the skull was gently removed using a sharp bent needle, followed by four stainless steel screws fixed from both sides of the craniotomy. Prelimbic cortex (PRL), infralimbic cortex (IL), ventral orbitofrontal cortex (vOFC), nucleus accumbens (shell - AcbS, and core - AcbC), and lateral septum (LS) were targeted with the EArs (see Table S2 for coordinates), with the AcbS electrode (A/P = +1 mm, L/M = ±2 mm, D/V = −7.6 mm) serving as a reference for the coordinates of other electrodes. The EAr electrode tips were dipped in DiI (1,1’-Dioctadecyl3,3,3’,3’-tetramethylindocarbocyanine perchlorate; 42364, SigmaAldrich) before implantation to mark their targets *post mortem*. Lastly, using toothpicks, a mixture of super-glue and dental cement was applied to the skull area surrounding the implant, to fix the implant to the skull. The animals were provided with painkiller (Meloxicam 1mg/kg) and antibiotic (Baytril, 5mg/kg) for three days post-surgery. Experiments were performed after at least 7 days of recovery.

### Experimental setup

Experimental arena was as previously described (Netser et al., 2019). Two Plexiglas triangular stimulus chambers with a metal mesh (1.0 x 2.0 cm holes) at their bottom were set in two randomly chosen opposite corners of the arena as previously described (John et al., 2023). The food chambers had a denser mesh (0.5 x 0.5 cm holes) at its bottom, such that the rats couldn’t consume the food while investigating it.

### Social vs. food paradigm (SVF)

Subject rats used only for behavioral analysis (Males, n = 15; Females, n = 14) were either group-housed (grouped) or singly housed (isolated) for 7 days prior to the experiment. The experiment began with two consecutive days, when the animal was placed in an arena with empty chambers for 15 minutes of habituation. From the third day onwards, subject animals were placed in an empty arena for 10 minutes, followed by 10-min habituation in the arena with empty chambers. Then, a social stimulus and standard chow-food (Envigo) were placed inside the chambers and a 5-min recording session (encounter) commenced. The subject rats then went through the same paradigm following a 24-hour food-deprivation (24h FD) on day 4, and 48-hour food-deprivation (48h FD) on day 5.

For the EAr-implanted rats (Grouped n = 13, Isolated n = 14), we used the same paradigm but started the recording session 5 minutes before the encounter, and defined this period, when only empty chambers occupied the arena, as baseline (as shown in Fig 1A). The re-grouped rats (n = 11) performed the SVF task twice, first in isolation and then one week after their return to group-housing (Please see Table S2 for details).

**Figure 1.**
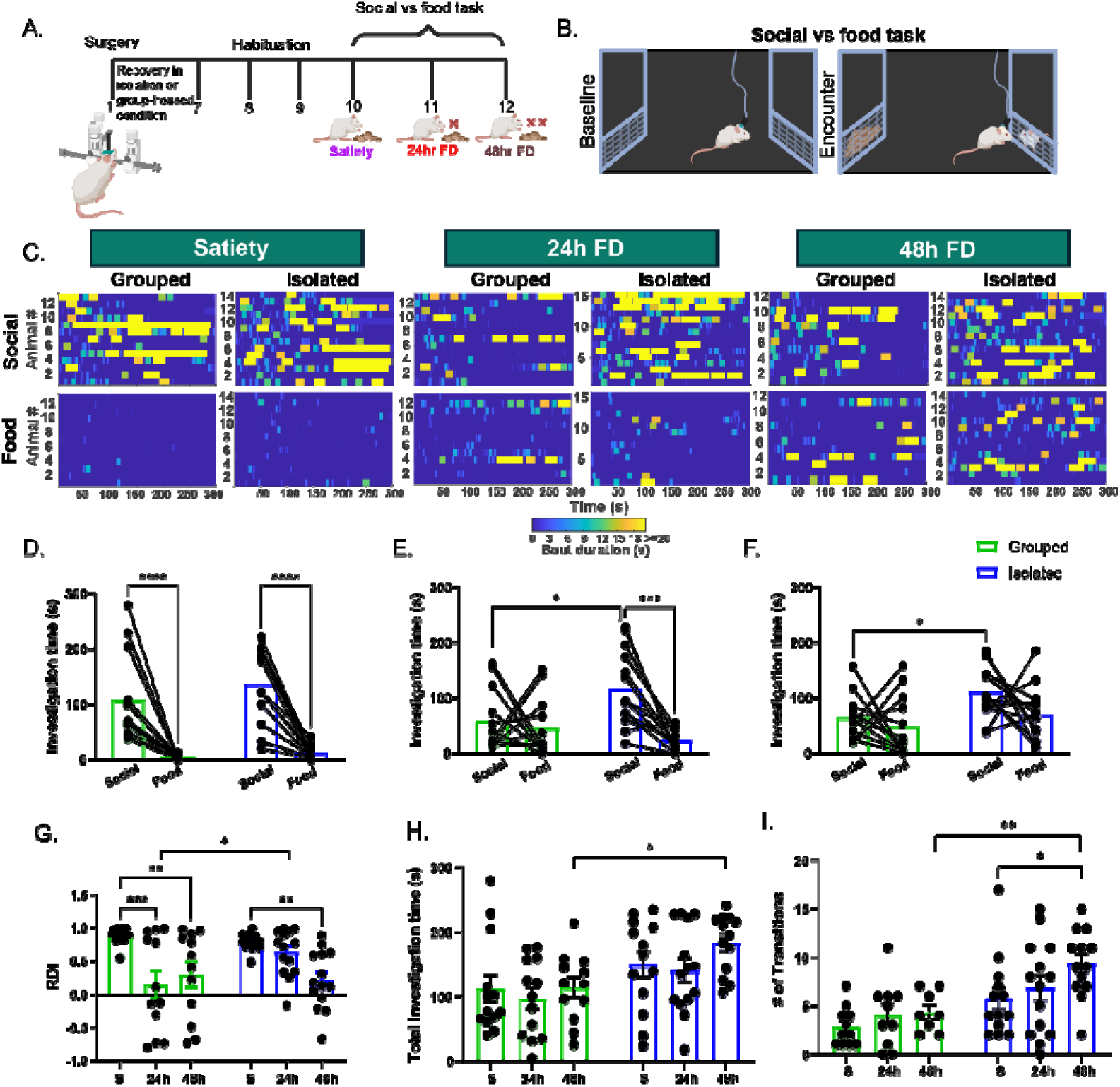
Behavioral differences between group-housed and socially isolated rats in social vs food task. **A**. A scheme of the experimental timeline (Created in https://BioRender.com). **B**. A scheme of the social vs food (SVF) task. C. Heatmaps of investigation bouts towards social or food stimuli during SVF task conducted by grouped and isolated rats at satiety (S), 24 hours of food-deprivation (24h FD, 24h), and 48 hours of food deprivation (48h FD, 48h). **D**. Statistical comparison between social and food investigation time for grouped and isolated rats at satiety (Two-way RM ANOVA, Stimulus: F = 62.65, P < 0.0001; Sidak’s MC test, Grouped, p < 0.0001, Isolated, p < 0.0001). E. Same as D, for 24h FD (Two-way RM ANOVA, Interaction between stimulus and housing: F = 5.69, p < 0.05, Stimulus: F = 10.4, P < 0.01; Sidak’s MC tests: social vs food in isolated, p < 0.001, grouped p > 0.1; grouped vs isolated in social, p < 0.05, food, p > 0.1). F. Same as D, for 48h FD (Two-way RM ANOVA, housing: F = 12.21, p < 0.01; Sidak’s MC test: grouped vs isolated in social, p < 0.05, food, p > 0.1). G-I. Comparison between grouped and isolated rats, for the relative discrimination index (RDI) in G (Two-way ANOVA, housing, F = 12.08, p < 0.001; Sidak’s MC test, grouped vs isolated at 48h FD, p < 0.05), number of transitions in H (Two-way ANOVA, housing, F = 16.01, p < 0.001; Sidak’s MC test, grouped vs isolated at 48h FD, p < 0.01), and total investigation time in I (Two-way ANOVA, interaction, F = 3.4, p < 0.05, housing, F = 11.9, p < 0.0001; Sidak’s MC test, grouped vs isolated at 24h FD, p < 0.05; Satiety vs 24h FD for grouped, p < 0.001, for isolated p > 0.1; Satiety vs 48h FD for grouped, p < 0.01, for isolated, p < 0.01). *p < 0.05, **p < 0.1, ***p < 0.001, ****p < 0.0001. All error bars represent SEM.

### Food vs. empty (FVE)

With some (n = 8) of the regrouped rats, we employed a FVE paradigm while recording their behavior and brain activity. In that case, 20-25 minutes after the SVF task the rats conducted a similar paradigm while presented with a chamber containing standard chow-food in one corner of the arena and an empty chamber in the opposite corner. This was done on each recording day, for all three metabolic conditions.

### Pharmacological manipulation

The orally-active Oxytocin Antagonist (OTA, (L-368,899) hydrochloride) was first diluted in sterile saline (1mg/mL). In rats used for behavioral analysis only, OTA solution (1mg/kg body weight; treatment group) or saline (control group) were intraperitoneally (IP) injected to the rats 45-50 minutes before the SVF session conducted after 24 hours of food-deprivation (day 2).

In EAr-implanted rats, SVF experiments were repeated with each rat twice (in randomized order): once with saline-injection after 24 hours of food-deprivation as a control, and another time with OTA injection, as described above. (see Table S2 for details).

### Electrophysiological recordings

As previously described (Mohapatra et al., 2022), following a brief (5-10 s) exposure to isoflurane, subjects were attached to the headstage of the electrophysiological recording system (RHD 32 ch, #C3314, Intan Technologies) through a custom-made Omnetics to Mill-Max adaptor (Mill-Max model 852-10- 100-10-001000). Electrophysiological signals were recorded via the RHD2000 evaluation system using an ultra-thin SPI interface cable, protected by an aluminium cylinder and connected a RHD2132 amplifier board (Intan Technologies). Recorded signals (sampled at 20 kHz) were synchronized with the video recording by a start signal sent through a custom-made triggering device and TTL signals from the camera to the recording system.

### Histology and electrode registration

After finishing all experiments, animals were sacrificed with an overdose of isoflurane and then perfused with 0.01 M phosphate buffer saline (PBS), followed by fixation in 4% paraformaldehyde (PFA) solution. After implant removal, brains were harvested and placed in PFA (4%) for 3-4 days, followed by sectioning of 50-um slices in the horizontal axis for electrode tracing using a vibratome (VT1000S, Leica). Sections were stained with DAPI and examined under a wide-field fluorescence microscope (Nikon Ti-eclipse) for verifying the placement of the electrode-tip marks (DiI fluorescence).

### Data analysis

#### Video tracking and behavioral analysis

Video clips were analysed using TrackRodent (algorithm WhiteRatsBodyBased for non-surgically implanted rats, WhiteRatsWiredHeadDirectionBased for surgically implanted rats) as previously described (Netser et al., 2019).

#### Electrophysiological analysis

All electrophysiological signals were analysed as previously described (Mohapatra et al., 2022). For the change in either theta (4-12 Hz) or gamma (30-80 Hz) power during the encounter stage, we subtracted the mean power averaged across the entire 5-min baseline stage from the one calculated for the 5-min encounter stage. We used the ‘mscohere’ function of MATLAB to estimate coherence values using Welch’s overlapped averaged periodogram method. The magnitude-squared coherence between two signals, x and y, was defined as follows:

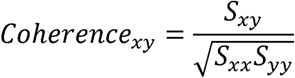

where S_xy_ is the cross-power spectral density of x and y; S_xx_ is the power spectral density of x and S_yy_ is the power spectral density of y. Coherence analysis was quantified only between brain regions pairs involved in at least five sessions of behavioral tasks.

#### Canonical Correlation Analysis (CCA)

CCA was used to quantify shared multivariate structure between behavioral and electrophysiological (LFP) features across experimental conditions. Prior to CCA, both datasets were pre-processed to ensure comparability across sessions. Electrophysiological features were standardized using *StandardScalar*() function from scikit-learn in python. Given a large number of features (28 features), we reduced the dimensions using principal component analysis (PCA), retaining enough components to explain 95% of the variance (number of PCA components determined individually for the dataset). Behavioral features were normalized using the *StandardScalar()* function without dimensionality reduction, as the number of behavioral variables was relatively small.

CCA was performed using the scikit-learn implementation (*CCA*()), with the number of canonical dimensions set to the smallest dimensionality of the two datasets (2 components in our analysis). To prevent overfitting, we implemented k-fold cross-validation (k = 5). In each fold, CCA was fit only on the training set, and canonical variates were computed for the test set. Final correlation estimates were averaged across folds.

To interpret the contribution of individual features, we computed structure coefficients for each modality, defined as the Pearson correlation between each original feature and its corresponding canonical variate (e.g., correlation of each electrophysiological feature with the neural canonical variate 1).

For group-level comparisons, pooled CCA was performed across all sessions regardless of their groups, followed by extraction of canonical scores (behavior canonical variates; LFP canonical variates) for each session. These scores were used for visualization of clustering patterns across food-deprivation and housing conditions. Statistical testing of canonical scores was performed using two-way ANOVA (Housing × Deprivation), followed by *post hoc* pairwise tests.

#### Network analysis

We constructed subject-level functional brain networks across four recorded brain regions (PRL, IL, AcbS, and AcbC) under each experimental condition. For each subject, coherence was computed separately for theta (4–12 Hz) and gamma (30–80 Hz) activity using DeepPhenotyping as previously described (Mohapatra et al., 2024).

Each connectivity measure yielded a weighted, undirected 4×4 adjacency matrix. To ensure consistency across methods, diagonal entries were set to zero and edge weights were normalized to the [0,1] range. Networks were computed separately for each frequency band.

#### Statistical analysis

Statistical tests were performed using GraphPad Prism version 9.0.0 for Windows, (GraphPad Software, San Diego, California USA, www.graphpad.com, see Table S1 for the details of all statistical analyses). data normality was tested using Kolmogorov-Smirnov or D’Agostino-Pearson omnibus (K2) test. Two-way ANOVA or Two-way repeated measures (RM) ANOVA was used to compare between multiple groups and variables. Upon finding a main effect or interaction in the ANOVA tests, Sidak’s or Dunn’s multiple comparison corrections were applied *post hoc*. Paired t-tests were used to compare the treatment groups.

All codes are available at https://github.com/98rishika/Social-vs-Food

## RESULTS

### Social isolation enhances social over food preference

To investigate how social isolation affects the balance between social and food-seeking motivations, we employed a social versus food (SVF) preference paradigm (Netser et al., 2020) in adult male rats chronically implanted with electrode arrays (EArs, see below), under varying conditions of food and social deprivation (Fig. 1A). Each session comprised a 5-min baseline stage, with subject animals exposed to two empty chambers in opposite corner of the experimental arena, followed by an encounter stage, with a social stimulus (a sex matched juvenile SD rat) occupying one chamber and food pellets in the other (Fig. 1B). Analysis of the subject’s investigation behavior (Fig. 1C) revealed that both group-housed (grouped) and singly housed (isolated, for 7 days prior to the experiment) rats exhibited a strong preference to investigate social stimuli over food when sated, with no significant difference between the two housing conditions (Fig. 1D). However, following 24 hours of food deprivation (24h FD), a clear divergence emerged between housing conditions. Group-housed rats displayed reduced social investigation time and showed no significant preference for either stimulus, consistent with an adaptive shift in their preference under metabolic need (Fig. 1E, green bars). In contrast, isolated rats maintained a significant social preference despite 24h FD and demonstrated significantly greater social investigation time compared to isolated rats in the same metabolic condition (Fig. 1E, purple bars). Moreover, even after 48 hours of food deprivation (48h FD), when both groups lost their social preference, isolated rats still exhibited elevated social engagement relative to group-housed controls (Fig. 1F). Accordingly, the relative discrimination index (RDI), reflecting the bias for social over food investigation, decreased significantly after 24h FD as compared to satiety in group-housed rats, but remained unchanged in isolated rats under the same metabolic condition (Fig. 1G). Moreover, isolated rats at 48h FD exhibited significantly higher total investigation time (Fig. 1H) and number of transitions between stimuli (Fig. 1I), compared to satiety, indicating generalized hyperactivity following social isolation.

Notably, all the behavioral patterns mentioned above were replicated in non-implanted male and female rats, confirming the robustness of the isolation-induced phenotype across genders (Fig. S1). Nevertheless, given their higher survival rate and superior recovery post-surgery, we conducted electrophysiological experiments with male rats only.

### Social isolation enhances theta and gamma oscillations in corticostriatal areas

To identify neural correlates of altered motivational hierarchies, we used the chronically-implanted EArs to record local field potential (LFPs) signals from six motivation-associated brain regions during the SVF task: prelimbic (PrL) and infralimbic (IL) areas of the mPFC, ventral OFC (vOFC), NAc core (AcbC) and shell (AcbS), and lateral septum (LS) (Fig. 2A-C).

**Figure 2.**
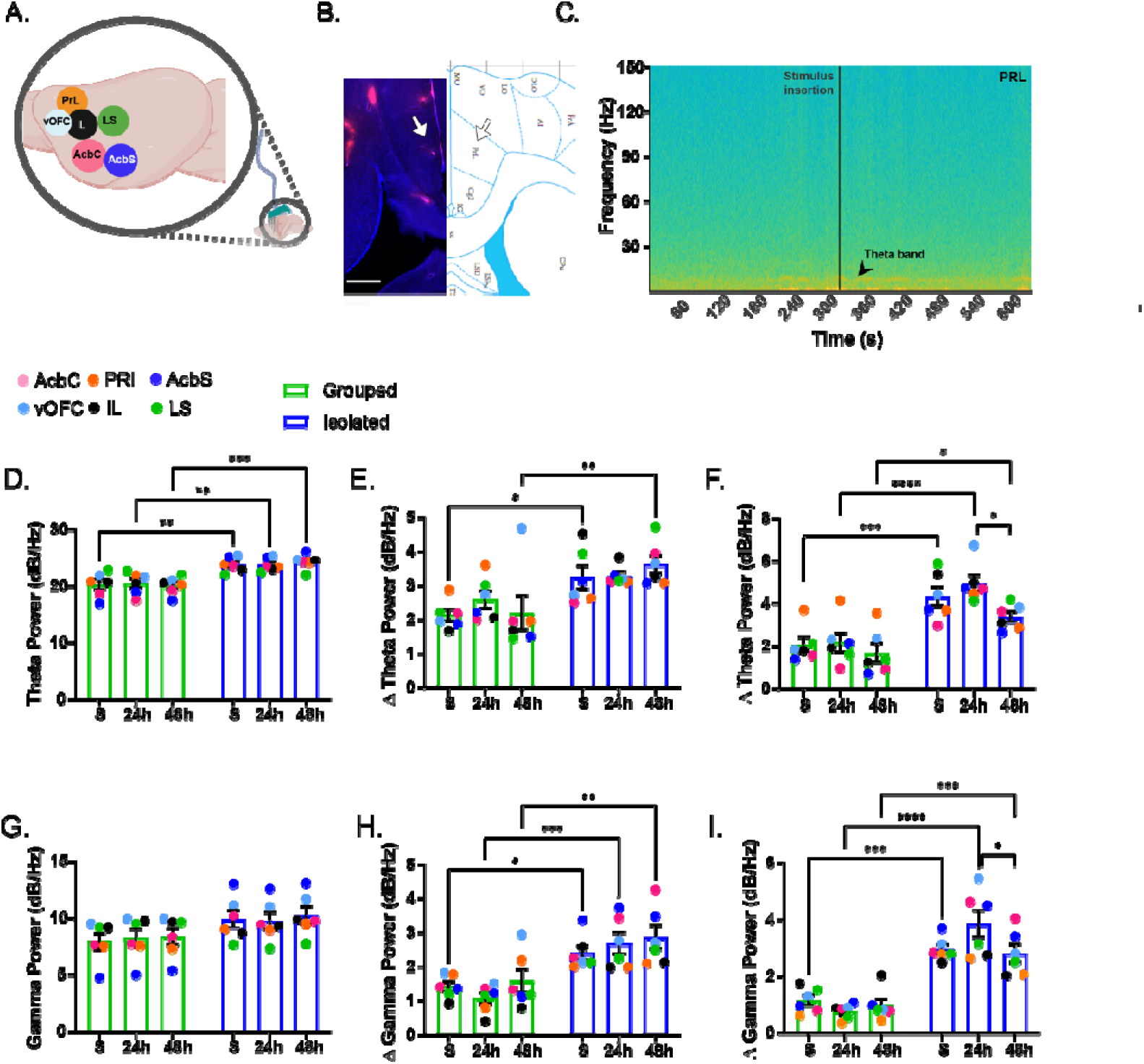
Isolated rats display higher induction of theta and gamma power during all stages of food-deprivation. **A**. A scheme of the targeted brain areas (Created in https://BioRender.com). **B**. A representative microscopy image (let) and rat brain atlas scheme (right), showing electrode tip tracing. Scalebar 1mm. C. An example spectrogram of LFP signals recorded from prelimbic cortex (PRL) from a rat during the baseline and encounter stages of the SVF task. **D.** Comparison of mean theta power during the baseline stage in grouped and isolated rats (Two-way ANOVA, housing conditions: F = 46.4, p < 0.0001; Sidak’s MC test, Grouped vs Isolated at satiety, p < 0.01, 24h FD, p < 0.01, 48h FD, p < 0.001). Each dot represents a brain area and is an average of values derived from 6-12 rats. **E**. Comparison of mean change in theta power during the whole encounter stage in the SVF task (Two-way ANOVA, housing conditions: F = 39.58, p < 0.0001; Sidak’s MC test, Grouped vs Isolated at satiety, p < 0.01, 24h FD, p < 0.05, 48h FD, p < 0.01). **F**. Comparison of mean change in theta power during social investigation bouts in the encounter stage of the SVF task (Two-way ANOVA, housing conditions: F = 46.4, p < 0.0001, Metabolic conditions: F = 3.669, p < 0.05; Sidak’s MC test: Grouped vs Isolated at satiety, p < 0.01, 24h FD, p < 0.01, 48h FD, p < 0.001; 24h FD vs 48h FD: Grouped, p > 0.1, Isolated, p < 0.05). G-I. Same as D-F for gamma power (G: Two-way ANOVA, housing conditions: F = 8.35, p < 0.01; H: Two-way ANOVA, housing conditions: F = 47.46, p < 0.0001; Sidak’s MC test, Grouped vs Isolated at satiety, p < 0.05, 24h FD, p < 0.001, 48h FD, p < 0.001; I: Two-way ANOVA, Interaction: F = 3.5, p < 0.05, housing conditions: F = 99.4, p < 0.0001; Sidak’s MC test, Grouped vs Isolated at satiety, p < 0.001, 24h FD, p < 0.0001, 48h FD, p < 0.001; 24h FD vs 48h FD for grouped, p > 0.1, Isolated, p < 0.05). *p < 0.05, **p < 0.1, ***p < 0.001, ****p < 0.0001. All error bars represent SEM.

Even during the baseline stage, before the introduction of social and food stimuli to the arena, isolated rats displayed significantly elevated theta power compared to grouped rats across all recorded regions (Fig. 2D). Moreover, upon stimulus presentation (encounter stage), isolated rats exhibited significantly greater increase in theta power than grouped rats across all metabolic states (except 24h FD, where only a trend was observed, Fig. 2E). This difference was even more significant during social investigation epochs, when theta power induction was highest in isolated rats in general, while showing a modest decrease at 48h FD (Fig. 2F). Similar results were found for LFP gamma power, although the difference in baseline gamma power between housing conditions did not reach statistical significance (Fig. 2G-I). These findings indicate that social isolation fundamentally alters LFP oscillatory dynamics in motivation-associated brain regions, especially during motivational competition between food and social stimuli.

### Re-grouping fails to reverse the isolation-induced behavioral and neural changes

To determine whether the effect of social isolation is reversible, we re-grouped isolated rats with their original cagemates for 7 days prior to behavioral testing. Unexpectedly, and unlike both isolated and group-housed animals, re-grouped rats continued to exhibit strong social preference even after 48h FD, thus presenting a new type of behavioral phenotype (Fig. 3A-B). To confirm that food deprivation was effective in re-grouped animals, we conducted a food versus empty chamber (FVE) task with re-grouped rats. These rats, which did not show any preference between the chambers at satiety, showed the expected increase in food investigation with increasing deprivation duration in the FVE paradigm (Fig. 3 C-D), thus confirming that motivation for food seeking according to the metabolic state is intact in re-grouped animals.

**Figure 3.**
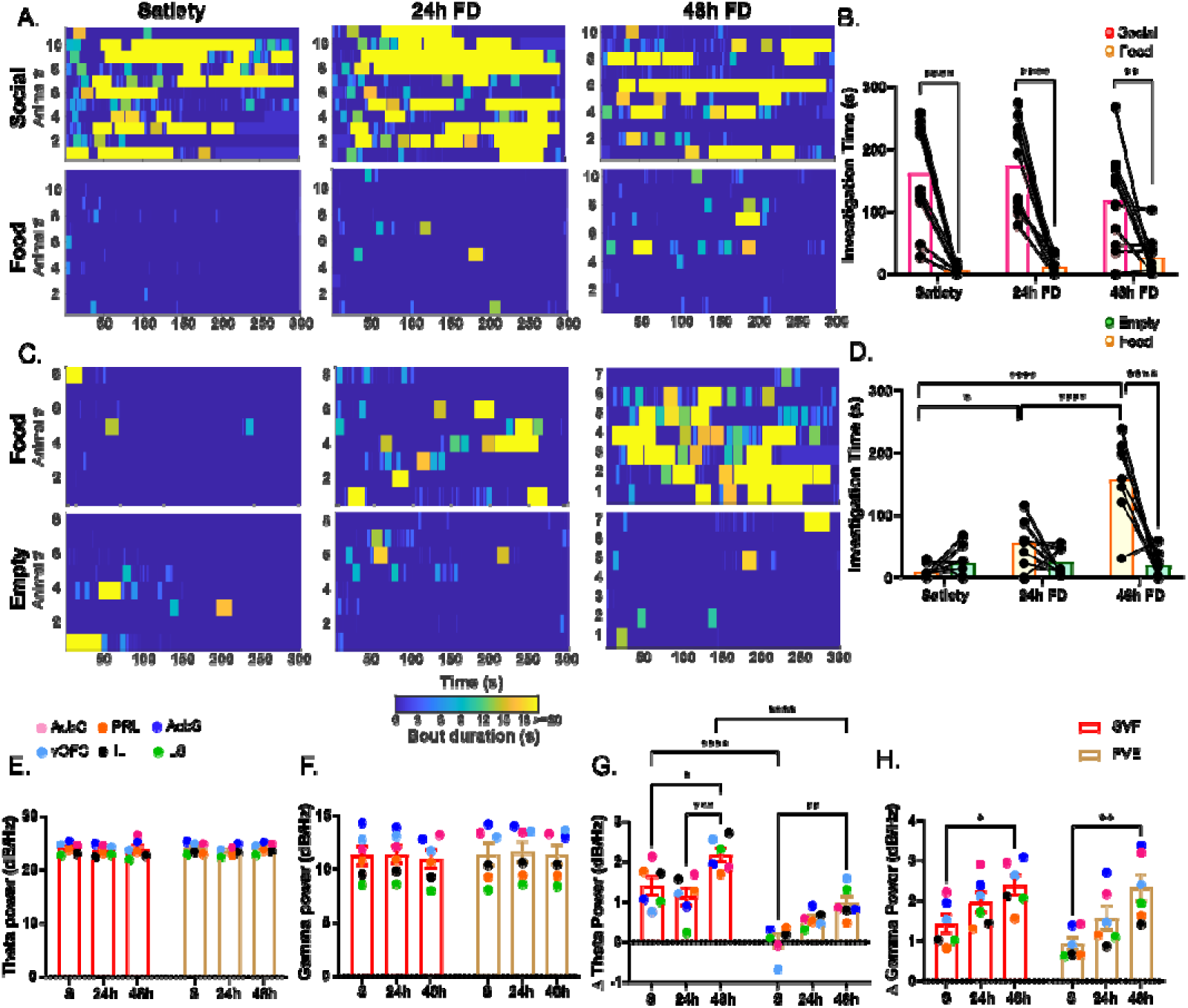
One-week re-grouping does not restore the behavioral phenotype of social isolation in rats. **A**. Heatmaps of investigation bouts towards social or food stimuli by re-grouped rats at satiety (S), 24 hours of food-deprivation (24h FD, 24h), and 48 hours of food deprivation (48h FD, 48h). **B**. Comparison of social vs food investigation time between satiety, 24h FD, and 48h FD in the re-grouped rats during the SVF task (Two-way ANOVA, Stimulus: F = 83.9, p < 0.0001; Sidak’s MC test, social vs food at satiety, p < 0.0001, at 24h FD, p < 0.0001, 48h FD, p < 0.01). **C-D**. Same as A-B, for food vs empty (FVE) task. (Two-way ANOVA, Stimulus: F = 17.62, p < 0.001, FD: F = 26.05, p < 0.0001, Interaction: F = 12.98, p < 0.001; Sidak’s MC test, food vs empty at 48h FD, p < 0.0001; Food at S vs 24h, p < 0.05, S vs 48h, p < 0.0001, 24h vs 48h, p < 0.0001). **E**. Comparison of mean theta power at the baseline stage between SVF and FVE tasks performed by re-grouped rats. F. Same as I, for gamma power. G. Comparison of mean change in theta power during the encounter stage between the SVF and FVE tasks (Two-way ANOVA, Metabolic conditions: F = 14.14, p < 0.0001, SVF vs FVE, p < 0.0001; Sidak’s MC test, satiety vs 48h FD during SVF, p < 0.05, FVE, p < 0.01; 24h FD vs 48h FD during SVF, p < 0.001, FVE, p > 0.1; SVF vs FVE at satiety, p < 0.0001, 24h FD, p = 0.07, 48h FD, p < 0.0001). H. Same as G, for gamma power (Two-way ANOVA, Metabolic conditions: F = 10.6, p < 0.001; Sidak’s MC test, Satiety vs 48h FD during SVF, p < 0.05, FVE, p < 0.01). *p < 0.05, **p < 0.1, ***p < 0.001, ****p < 0.0001. All error bars represent SEM.

Neural recordings from re-grouped rats revealed complex interactions between housing history and metabolic state. As expected, no difference was observed during baseline between the SVF and FVE tasks, as well as between the metabolic conditions, for both theta and gamma power. Moreover, both theta and gamma power induction during the test increased substantially after 48h FD for both tasks (Fig. 3E-H). Nevertheless, the induction of theta power (but not gamma power) during the test was significantly more robust for the SVF task as compared to FVE, strongly suggesting that even after 48 hours of FD most induction of theta power was associated with the presence of a social stimulus rather than food (Fig. 3G-H). Notably, during the SVF task, re-grouped rats showed theta power changes resembling group-housed rats at satiety and 24h FD, but exhibited a significant increase at 48h FD, suggesting a complex interaction between metabolic pressure and housing history, leading to persistent isolation-induced neural adaptations (Fig. 3G; also see Fig S2 B-C). Together, these results indicate that a 1-week re-socialization is insufficient to reverse the behavioral and neural consequences of social isolation. Instead, re-grouped animals seem to create a class of their own, from both the behavioral and neural aspects.

### Social isolation increases corticostriatal network coherence during social versus food decision-making

To examine how social isolation affects coordinated neural activity across the recorded network of motivation-associated brain regions, we analyzed the coherence between LFP signals simultaneously recorded across the various brain areas. Our analysis focused on four regions (PRL, IL, AcbC, and AcbS), constructing six pairs of brain regions with sufficient sample size (n≥5 rats) across all experimental groups. During baseline, theta and gamma coherence did not differ significantly between housing conditions (Fig. S2 D-E). However, during the encounter stage of the SVF task, we found significant differences between them. Group-housed rats showed minimal changes in network coherence across food-deprivation states, with consistently low coherence change in both theta and gamma bands (Fig. 4A-D). In contrast, isolated rats displayed significantly elevated theta coherence across the corticostriatal network. Specifically, during satiety isolated animals showed theta coherence change which was significantly higher from both grouped and re-grouped animals (Fig. 4A). As demonstrated by the theta coherence network analysis displayed in Fig. 4B, this was most clearly reflected in the strength of PRL as a node in this network (note the larger size of the PRL orange node in Fig 4B; See legend and table S for stats). This result suggests that the PRL could be playing an important part in maintaining the functional coordination within this circuit based on the housing condition. Gamma coherence also increased in isolated rats, although the effect was less pronounced than for theta coherence (Fig. 4C). Re-grouped rats showed intermediate coherence levels, with patterns that partially resembled isolated rats, particularly at 48h FD (Fig. 4D). Altogether, these findings demonstrate that social isolation enhances functional coordination within corticostriatal circuits, potentially reflecting altered information processing during a motivational conflict.

**Figure 4.**
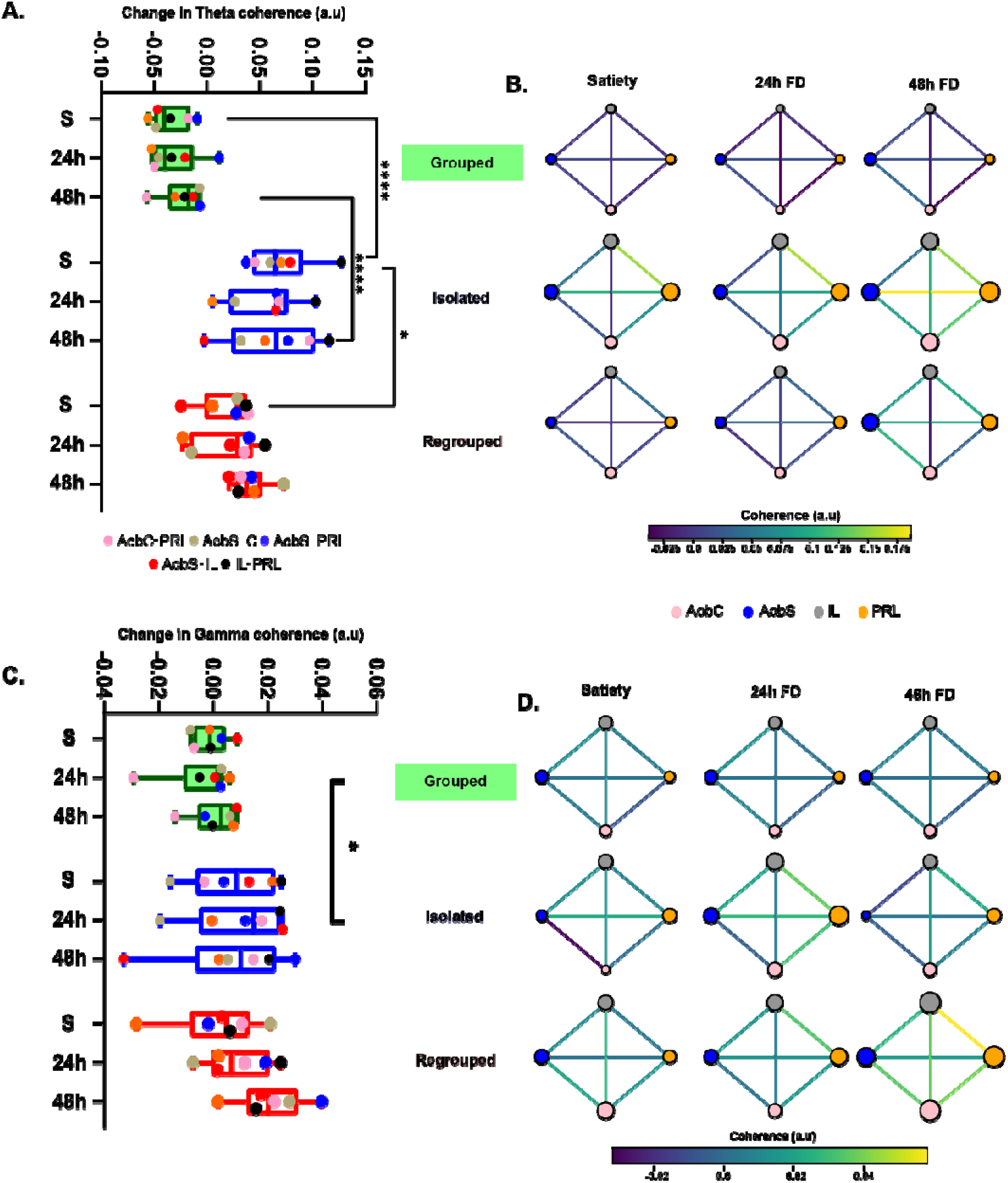
The corticostriatal network exhibits higher coordination during the SVF task in isolated rats. **A.** Comparison of mean change in theta coherence between pairs of brain areas recorded in grouped, isolated, and re-grouped animals (Two-way ANOVA, housing conditions: F = 47.8, p < 0.0001; Sidak’s MC test, Grouped vs Isolated at satiety, 24h FD and 48h FD, p < 0.0001; Grouped vs regrouped at satiety and 24h FD, p < 0.05, 48h FD, p < 0.01; Isolated vs regrouped at satiety, p < 0.05). Each dot represents a pair of brain regions and is an average value derived from at least 5 rats. **B.** Network graphs obtained from the theta coherence values in the three housing conditions and three metabolic states (Node Strength of PRL, Two-way ANOVA, housing conditions: F = 3.276, p < 0.05, Sidak’s MC test, Grouped vs isolated, p < 0.05). C-D. Same as A-B, for gamma coherence (Two-way ANOVA, housing conditions: F = 3.26, p < 0.05). *p < 0.05, **p < 0.1, ***p < 0.001, ****p < 0.0001. All error bars represent SEM.

### Canonical correlation analysis reveals dissociated neural signatures of metabolic state and housing condition

To quantify the multivariate relationships between the behavioral and neural features, we performed canonical correlation analysis (CCA) (Hotelling, 1992). CCA is a statistical tool analysing the relationship between two sets of multivariate variables derived from the same subjects. It aims to find linear combination of variables, called canonical variates, that maximize the correlation between the two sets. In our study, the first canonical space captured shared variance primarily related to metabolic state, such that sessions were clearly segregated based on whether rats were sated or food-deprived, regardless of housing condition (Fig. 5A-C). This segregation was evident in the behavioral canonical axes, as shown in the heatmaps. The sessions with more social investigation are segregated on the right side of the canonical space, whereas the ones with more food investigation are towards the left. Statistical analysis of the first LFP-derived canonical variate confirmed significant main effects of metabolic state across all housing groups, with many *post hoc* significant differences between the various metabolic states (Fig. 5D). Most specifically, we found a significant (or borderline significant) difference in this canonical dimension between satiety and 48 h FD across all housing conditions. These results indicate that metabolic signals are robustly encoded by distributed neural activity patterns.

**Figure 5.**
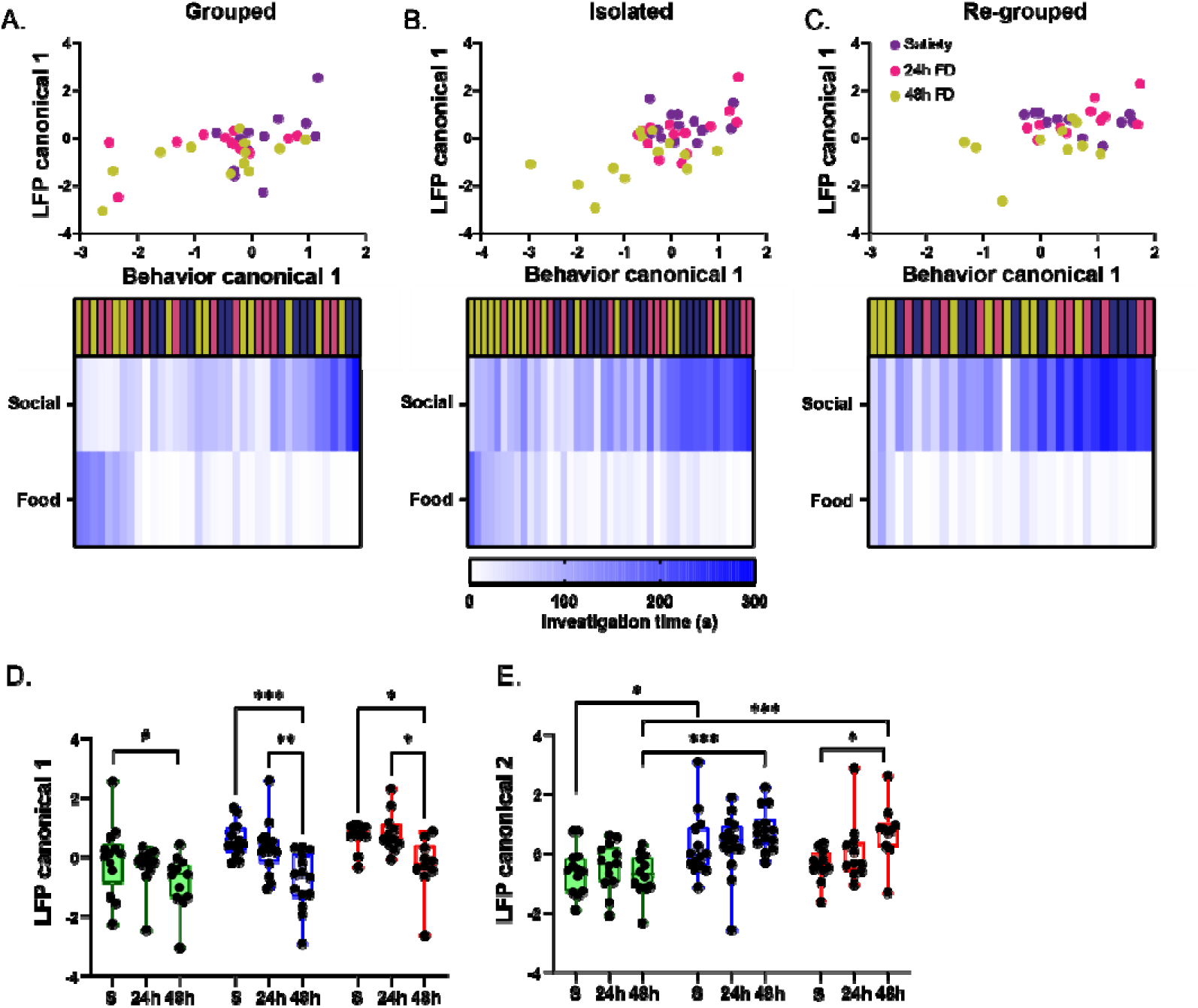
CCA reveals distinct canonical neural-behavioral spaces distinguishing between metabolic and housing conditions. **A.** Distribution of grouped rats in first canonical space derived from the CCA. **B**. Same as A, for isolated rats. **C**. Same as A, for re-grouped rats. The heatmaps display social and food investigation time of each session (satiety in magenta, 24h FD in pink, and 48h FD in golden) arranged in increasing order of the behavioral canonical variate. Notice that the sessions with more social investigation are towards the right, and the ones with food investigation are towards the left. **D**. Comparison of the LFP-derived canonical variates of the first canonical space across housing and metabolic conditions (Two-way ANOVA, Housing conditions: F = 16.07, p < 0.0001, Metabolic conditions, F = 8.5, p < 0.001; Sidak’s MC test, Satiety vs 48 h FD for grouped, p = 0.07, Isolated, p < 0.001, Regrouped, p < 0.05; 24h FD vs 48h FD for grouped, p > 0.1, Isolated, p < 0.01, Regrouped, p < 0.05). E. Same as D for the second canonical space (Two-way ANOVA, Housing conditions: F = 13.9, p < 0.0001, Metabolic conditions, F = 3.09, p < 0.05; Satiety vs 48h FD for grouped, p > 0.1, Isolated, p > 0.1, Regrouped, p < 0.05; Grouped vs Isolated at satiety, p < 0.05, 24h FD, p > 0.1, 48h FD, p < 0.001; grouped vs regrouped at satiety and 24h FD, p > 0.1, 48h FD, p < 0.001). *p < 0.05, **p < 0.1, ***p < 0.001, ****p < 0.0001. All error bars represent SEM.

In contrast to the first variate, the second LFP canonical variate differed significantly between housing groups, with the largest separation observed between grouped and isolated rats at 48h FD (Fig. 5E). Notably, in most cases the metabolic states did not differ from each other for a given housing condition. Together, these results demonstrate that these two canonical dimensions encode orthogonal sources of variance: one primarily reflecting metabolic state (physiological drive) while the other reflecting housing condition (social drive). This dissociation highlights the existence of separate neural-behavioral coupling patterns for these two fundamental determinants of motivated behavior.

### Oxytocin receptor antagonism reverses isolation-induced enhancement in social preference

Given the established role of oxytocin in regulating both social and appetitive behavior, we hypothesized that manipulating oxytocin signalling might affect the isolation-induced enhancement of social over food preference. To test this hypothesis, we first injected either saline or an orally-active oxytocin receptor antagonist (OTA) to isolated rats 45-50 minutes prior to the SVF task conducted following 24h FD, without electrophysiological recordings. Similar to untreated isolated rats (Fig. 1F), saline-treated isolated rats kept exhibiting a significant social preference even at 24 h of FD (Fig. 6A-E, Fig. S3A-D). In contrast, OTA-treated isolated rats showed balanced investigation of both social and food stimuli, with no significant preference for any of them (Fig. 6E), thus resembling the behavior of untreated grouped rats under similar metabolic pressure (Fig. 1E). Accordingly, while saline-treated rats exhibited no change in RDI between satiety and 24 h FD, OTA-treated rats showed a significant reduction in their RDI at 24 h FD, reflecting the loss of their social over food preference (Fig. 6I, S3E). This occurred without significant changes in total investigation time or number of transitions between stimuli (Fig. 6G-H), suggesting no effect of OTA treatment on motor activity or general arousal of the animals. Altogether, these findings establish that oxytocin receptor signalling is necessary for the isolation-induced change in motivational hierarchies.

**Figure 6.**
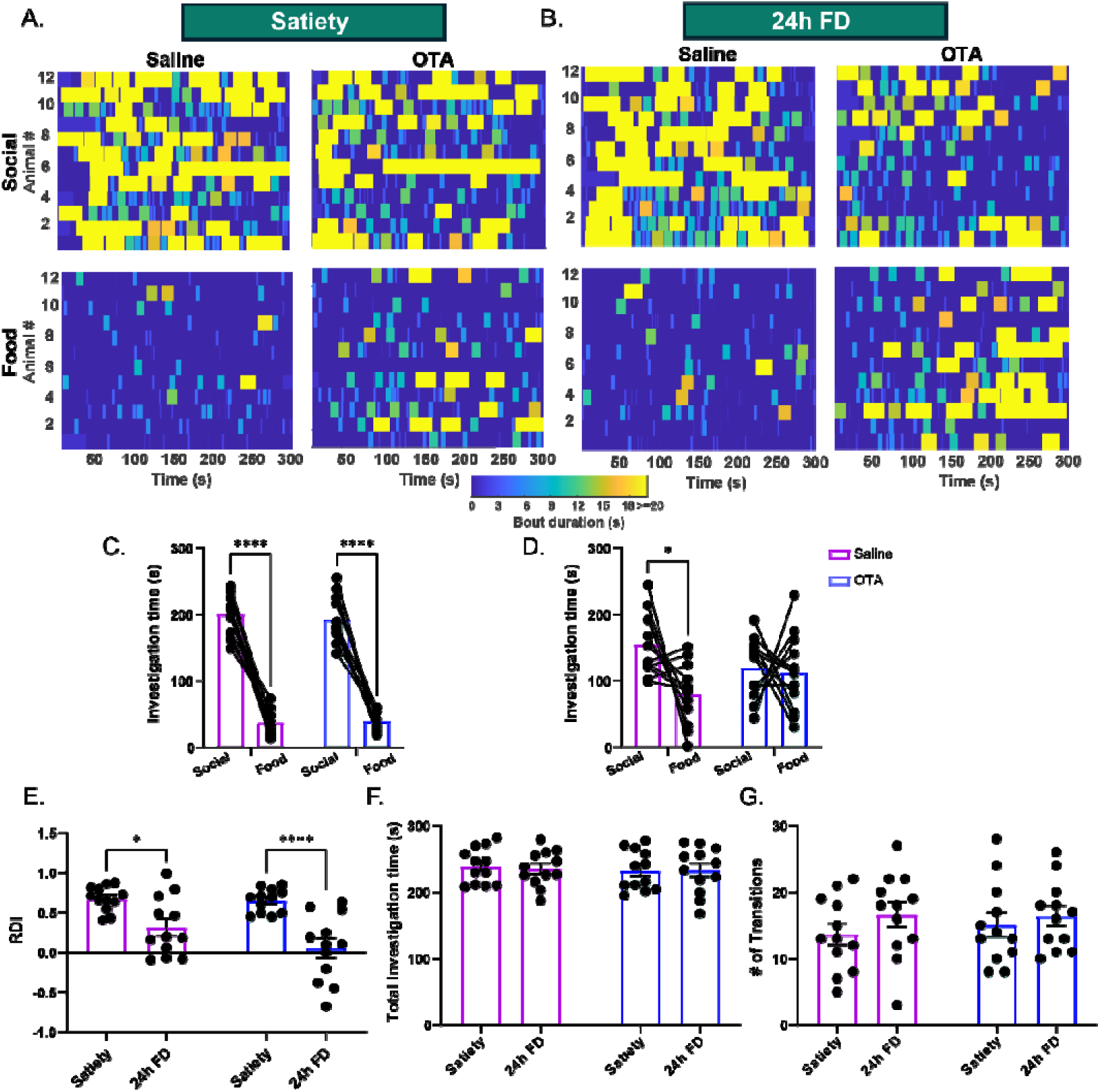
Treatment with oxytocin receptor antagonist blocks the enhancement of social over food preference isolated rats. **A-B**. Heatmaps of investigation bouts towards social or food stimuli between saline and oxytocin receptor antagonist (OTA)-treated rats at satiety (A) and 24 hours of food-deprivation (24h FD) (B) **C**. Comparison of the mean social and food investigation times between saline and OTA-treated rats at satiety (Two-way ANOVA, Stimulus, F = 305.6, p < 0.0001; Sidak’s MC test, social vs food in saline group, p < 0.0001, treatment group, p < 0.0001). **D**. Same as C, for 24h FD (Two-way ANOVA, Stimulus, F = 4.579, p < 0.05; Sidak’s MC test, social vs food in saline group, p < 0.05, treatment group, p >0.1). **E-G.** Comparison between saline and OTA-treated rats of the relative differential index (RDI) in E (Two-way ANOVA, Treatment, F = 30.5, p < 0.0001; Satiety vs 24h FD for saline group, p < 0.05, treatment group, p < 0.0001; Saline vs OTA at 24h FD, p = 0.07),, total investigation time in F, number of transitions in G. *p < 0.05, **p < 0.1, ***p < 0.001, ****p < 0.0001. All error bars represent SEM.

### Oxytocin receptor antagonism selectively reduces theta power in corticostriatal regions

To reveal neural correlates of the OTA behavioral effect, we applied the same protocol as described above to Ear-implanted animals. These animals repeated the same behavior as described above for unrecorded animals, although the social over food preference of saline-treated animals was just borderline significant, most probably because they were tethered to the electrophysiological recording system (Fig. S3A-E). We then compared LFP oscillations in OTA- versus saline-treated isolated rats. The experimental design included within-subject controls: each rat underwent the SVF paradigm twice over consecutive weeks at satiety and under 24h FD, receiving saline in one week and OTA in the other (order randomized; saline-first: n=4, OTA-first: n=5; See Table S2 for details). Because pilot data from group-housed rats revealed significant attenuation of theta power induction during repeated testing (Fig. S4A-B), we normalized the 24h FD data to each rat’s baseline recorded at satiety, to account for potential order effects.

Baseline theta and gamma power did not differ between saline- and OTA-treated groups (Fig. 7A, D). However, OTA treatment significantly reduced theta power induction during the entire encounter stage at 24h FD, compared to saline controls (Fig. 7B). This suppression was even more pronounced when analysing theta power specifically during social investigation bouts (Fig. 7C). Interestingly, OTA treatment at 24h FD increased gamma power during social investigation bouts (Fig. 7F), suggesting a dissociation between theta and gamma power under OTA treatment.

**Figure 7.**
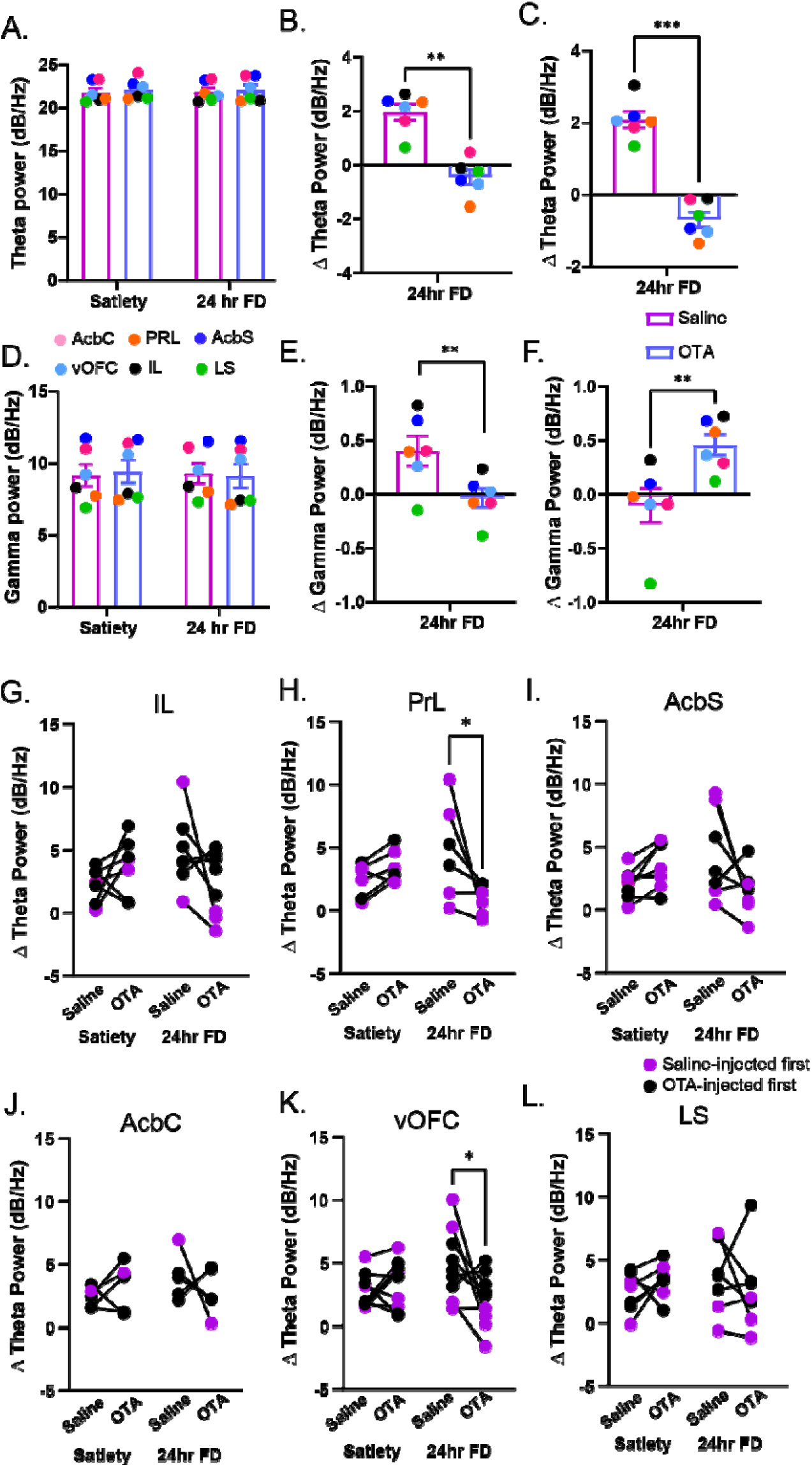
Systemic OTA treatment decreases theta power induction during SVF task. **A.** Comparison of mean theta power during the baseline stage of the SVF task, in saline and OTA-treated rats. Each dot represents a brain area and is an average of values derived from 5-9 rats. **B**. Comparison of mean change in theta power during the encounter stage of the SVF task, between saline and OTA-treated rats (Paired t-test, p < 0.01). C. Comparison of mean change in theta power during social investigation bouts during the SVF encounter stage between saline and OTA-treated rats (Paired t-test, p < 0.001). **D-F**. Same as A-C, for gamma power. E. (Paired t-test, p < 0.01). F. (Paired t-test, p < 0.01). **G-L**. Comparison of the change in theta power in the individual brain areas. H. Two-way RM ANOVA, Interaction, F = 8.0, p < 0.05; Saline vs OTA at 24h FD, p < 0.05. I. Two-way RM ANOVA, Interaction, F = 6.99, p < 0.05; Saline vs OTA at 24h FD, p = 0.057. K. Two-way RM ANOVA, Interaction, F = 5.7, p < 0.05; Saline vs OTA at 24h FD, p < 0.05. *p < 0.05, **p < 0.1, ***p < 0.001. All error bars represent SEM.

Finally, region-specific analysis revealed that the OTA-induced theta suppression was statistically significant specifically in PRL and vOFC (Fig. 7 G-L). These findings suggest that oxytocin receptor signalling regulates motivational hierarchies through brain region-selective modulation of theta oscillations.

## Discussion

This study provides insights into neural mechanisms that arbitrate between internal states. The results of this study should be interpret in light of the social homeostasis hypothesis, a framework proposing that organisms actively detect and regulate the quantity and quality of social contact to maintain an optimal set-point (Lee et al., 2021; Matthews & Tye, 2019). A key prediction of social homeostasis hypothesis is that social deprivation triggers compensatory behavioral and neural responses to restore social contact. However, a critical unresolved question is how social homeostasis interacts with other fundamental homeostatic drives, particularly when competing needs are simultaneously active.

Our results show very clearly a competition between the social and metabolic needs. On one hand, SD rats show a clear social over food preference during satiety but lose this preference upon increase of the metabolic need, after 24 hours of food deprivation. On the other hand, under heightened social need following social isolation, they exhibit social over food preference even after 24 hours of food deprivation. Thus, deprivation of either food or social interactions tip the preference scale in the direction of deprivation compensation. In light of the social homeostasis theory (Matthews and Tye, 2019), these results may suggest that both the social control and energy control centers are equally competing on the same activator system. Nevertheless, our observation that re-grouped rats maintain enhanced social preference even under food deprivation reveals a long-term reorganization in the hierarchy of competing motivational drives.

Isolated rats exhibited elevated theta power across all recorded brain regions during both baseline and encounter stages. Theta oscillations in corticostriatal circuits have been implicated in reward anticipation, approach behavior, and coordination of distributed neural ensembles during motivated states (Donnelly et al., 2014; Herweg et al., 2020; Mohapatra et al., 2024, 2025). The heightened theta power in isolated rats may reflect increased arousal, enhanced attention to rewarding stimuli, or altered synchronization of neural populations encoding motivational salience. Notably, theta power in re-grouped animals was significantly higher in the SVF task compared to the FVE paradigm, in both satiety and food deprivation conditions. This indicates that, at least in re-grouped animals, the effect of the presence of social stimuli on theta rhythmicity in the recorded areas is much stronger than that of food stimuli, even under food deprivation condition. This result may explain why re-grouped rats still show social over food preference even under 24 h of food deprivation.

Our results are in accordance with a recent publication (Rocha-Almeida et al., 2025) which used a similar recording technique to investigate corticostriatal LFP power changes during a competition between food and social interactions in Lister Hooded rats, a rat model of ADHD (Jogamoto et al., 2020). Interestingly, these animals, which showed a clear preference for food over social interactions even in satiety, also showed a reduction in theta power before social interaction, in the same brain regions recorded by us. These results further support our conclusion that theta rhythmicity in corticostriatal circuits is strongly associated with social contexts.

Gamma oscillations, which are associated with local circuit processing and sensory integration (Buzsáki & Wang, 2012; Jensen et al., 2007), were also elevated in isolated rats during the encounter stage. The concurrent enhancement of both theta and gamma power suggests that social isolation affects multiple temporal scales of neural computation, potentially altering both long-range coordination and local processing efficiency within reward circuits.

A central finding of our study is that social isolation dramatically enhances functional coupling within corticostriatal circuits during motivational decision-making, as isolated rats exhibited markedly elevated theta coherence across the recorded network. This enhanced theta coherence persisted across all food-deprivation states and remained partially elevated even after one week of re-grouping, further indicating a long-term reorganization of network dynamics. The mPFC plays a critical role in representing internal states and integrating them with external sensory information to guide goal-directed behavior (Bein & Niv, 2025; Mohapatra & Wagner, 2023), while the NAc encodes reward value and motivation (Marinescu & Labouesse, 2024). Increased synchronization between these regions may create a neural state that preferentially channels social information through reward circuits, biasing behavioral output toward social approach even when metabolic signals would normally redirect attention to food. The intermediate coherence levels observed in re-grouped rats are particularly informative for understanding the temporal dynamics of social homeostatic regulation. Despite one week of group-housing, re-grouped animals maintained elevated coherence compared to group-housed controls, especially under severe food deprivation (48h FD). In line with the behavioral results, these findings suggest that social isolation does not merely enhance social motivation but rather reorganize the internal hierarchy between different homeostatic needs. In the framework of social homeostasis, this can be interpreted as a compensatory adjustment of the social homeostatic set-point, which takes precedence over metabolic homeostatic signals when social needs are unmet for a longer time. Thus, changes in network coherence may reflect modified integration of social and metabolic information, making their combination more sensitive to prior social experience.

Our canonical correlation analysis (CCA) revealed a critical insight: metabolic state and housing condition are encoded in different orthogonal neural dimensions. The first canonical space primarily captured variance related to food deprivation, successfully segregating sated from food-deprived animals regardless of housing condition. This demonstrates that the recorded neural circuits significantly track metabolic signals even during social isolation. In contrast, the second neural canonical variate distinguished housing conditions. This dissociation suggests that social isolation does not disrupt the encoding of metabolic state but rather creates an independent neural signature. The fact that these two dimensions are orthogonal suggests that social isolation-induced changes involve distinct activity patterns from those encoding metabolic states.

It was previously shown that the level of oxytocin in plasma of the rats is enhanced by social isolation (Neal et al., 2018) and that systemic administration of OTA induces a decrease in social interaction after social isolation in mice (Liu et al., 2025). In our study, treatment with OTA restored balanced investigation of social and food stimuli following 24h FD in isolated rats, effectively normalizing their behavior to resemble the behavior of grouped animals. Importantly, this behavioral normalization occurred without affecting general motor activity or arousal, indicating specificity for motivational processing rather than non-specific activation. The neural mechanism which accompanied this behavioral change involved selective suppression of theta oscillations, particularly in the PRL and vOFC. This regional specificity is notable, given that among the recorded structures PRL, IL and LS show high oxytocin receptor expression, while NAc and OFC exhibit sparse receptor density (Ostrowski, 1998). The observed theta suppression in regions with varying receptor expression suggests that oxytocin may exert its effects through direct modulation of oxytocin receptor highly expressing regions (e.g., PRL), which then influence downstream targets through network interactions. While theta power was strongly suppressed, gamma power during social investigation increased following OTA treatment. This differential effect of OTA administration suggests that oxytocin receptor signaling specifically regulates slower oscillatory rhythms that may coordinate information across distant brain regions, while leaving local gamma-band processing relatively intact or even disinhibited. The dissociation of OTA effects on theta and gamma oscillations is in accordance with the fact that theta, but not gamma power was different between the SVF and FVE paradigm under the same metabolic conditions. Together, these observations further support the idea that theta oscillations in the corticostriatal system is strongly and specifically affected by the social context.

A recent work (Rea et al., 2025) demonstrated that oxytocin signaling in the dorsal hippocampus specifically enhances food intake in the presence of familiar conspecifics while having no effect during isolated feeding. Importantly, knockdown of hippocampal oxytocin receptors eliminated both this social facilitation of eating and impaired social recognition memory, suggesting that oxytocin coordinates social context with feeding behavior across distributed neural circuits. While this study examined how social context modulates feeding through hippocampal oxytocin signaling, our results reveal a complementary mechanism: corticostriatal oxytocin signaling determines motivational priorities when social and food rewards compete directly. Together, these findings suggest that oxytocin operates as a multi-regional integrator of social and metabolic information, with circuit-specific roles ranging from hippocampal encoding of social context during feeding to corticostriatal regulation of motivational hierarchies during reward competition.

Limitations of the study: Several limitations should be noted. We used systemic OTA administration. Thus, the effect of it on the recorded signals may be indirect and not necessarily involving oxytocin receptors in the recorded areas. While we identified regional differences in theta suppression following OTA treatment, the specific cell types and circuit mechanisms remain to be determined. Also, how the duration and timing of isolation effects on neural plasticity warrant further investigation. Moreover, we revealed that one week of re-grouping does not fully reverse the effect of acute social isolation. Yet, it is still not clear if that effect is completely irreversible or does it demand more re-grouping time to be reversed. Finally, investigating how social isolation affects other forms of motivated behavior beyond social interaction and food seeking would provide broader insights into how the brain arbitrates between concurrent needs to guide behavior.

### Summary

Our findings address several key unresolved questions in the social homeostasis framework. First, they demonstrate that the social homeostatic system does not operate independently but rather must continuously negotiate with other fundamental homeostatic drives. The ability of social isolation to override metabolic signals even after re-grouping reveals the potency of social homeostatic regulation and suggests that, at least in SD rats, social needs may occupy a hierarchically privileged position relative to other needs in some circumstances.

Second, our results establish neural theta power and coherence in the corticostriatal system as a measurable biomarker of heighten social need state. The graded increase in theta coherence with isolation duration and its partial persistence after re-grouping suggest that coherence measures could quantify the degree of social homeostatic imbalance and track recovery dynamics.

Third, the identification of oxytocin-regulated theta oscillations as a potential mechanism provides a specific neural substrate for social homeostatic regulation by neuromodulators. This finding bridges between distinct levels of analysis, from molecular signalling (oxytocin receptors) to systems-level neural dynamics (theta oscillations) and behavior (motivational choice), offering a relatively comprehensive account of how social experience shapes motivational hierarchies.

## Supporting information

Table S1 - Stats

Table S2 - Summary of experiments

## Acknowledgements

We acknowledge the use of Claude (Opus 4.5) to improve the language, clarity, and academic tone of this manuscript. All AI-generated suggestions were reviewed and edited by the authors, who take full responsibility for the final content. We thank Alex Bizer, the experimental systems engineer of the Faculty of Natural Sciences of the University of Haifa for technical help.

## Funding

This study was supported by the Israel Science Foundation (2220/22 to SW), the Ministry of Health of Israel (3-19884 for PainSociOT to SW), the German Research Foundation (DFG) (SH 752/2-1 to SW), the United States-Israel Binational Science Foundation (2019186 to SW) and the European Research Council (ERC-SyG oxytocINspace).

## Author contributions

R.T.: Formal Analysis, Investigation, Methodology, Validation, Visualization, Writing - original draft, and Writing - review & editing; S.R.: Investigation, Analysis; S.N.: Data curation, Project administration, Software, Validation, Writing - original draft, and Writing - review & editing. S.W.: Conceptualization, Funding acquisition, Project administration, Resources, Supervision, Writing - original draft, and Writing - review & editing.

## Competing interests

The authors declare that they have no competing interests.

## Supplemental Information

**Table S1.** An Excel file containing details of all statistical analyses, organized by figure.

**Table S2.** An Excel file containing summary of experiments.

## Notes

https://github.com/98rishika/Social-vs-Food

### Competing Interest Statement

The authors have declared no competing interest.

### Summary of Updates

The order of figures and text was revised. Title was changed.

