## Supplementary material for "Dynamic corticostriatal neural coordination governs competition between social and metabolic drives in rats": Document S1

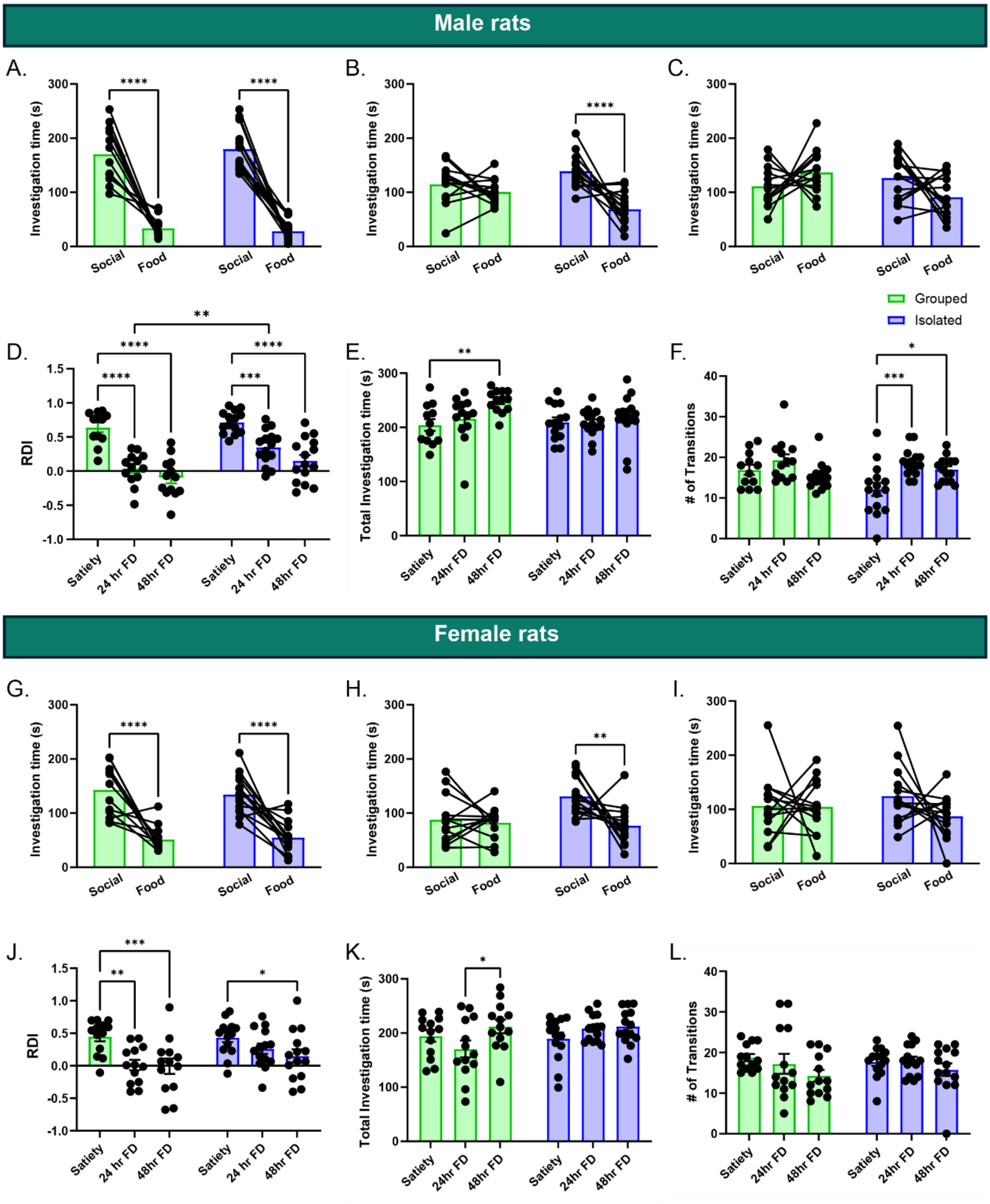


**Fig S1. Behavioral differences between group-housed and socially isolated non-implanted rats in social vs food tasks. A.** Statistical comparison between the social and food investigation time between grouped and isolated male rats at satiety. **B.** Same as F for 24h FD. **C.** Same as F for 48h FD. **D-F.** Comparison of the total investigation time in D, number of transitions in E, and relative differential index (RDI) in F, between grouped and isolated rats. **G-L.** Same as A-F for female rats.

*p<0.05, **p<0.01, ***p<0.001, ****p < 0.0001 post-hoc Sidak's multiple comparisons following detection of main effects by Two-way ANOVA.

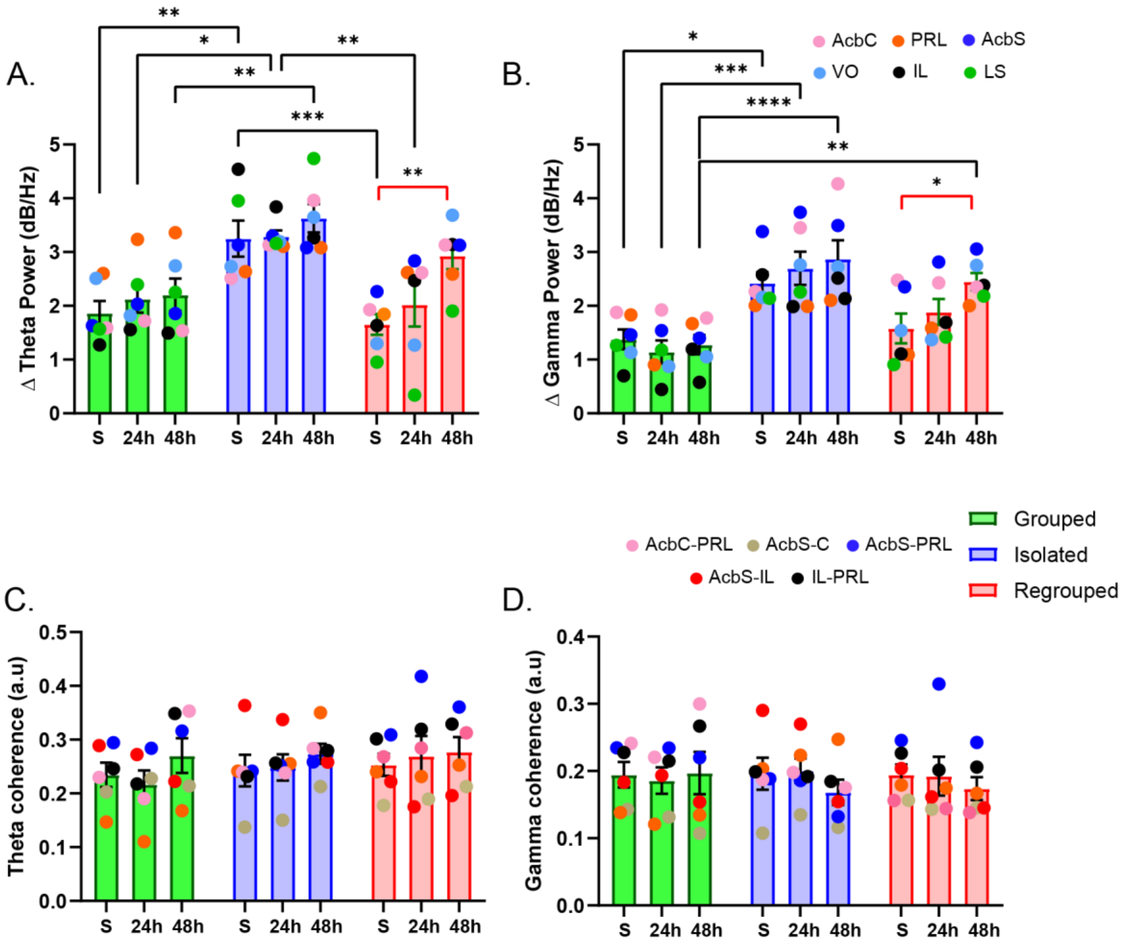


**Fig S2**. **A.** Comparison of change in theta power between Grouped, Isolated, and Regrouped rats on different days of food-deprivation. **B.** Same as A for gamma power. **C.** Comparison of theta coherence across six pairs of brain regions during the baseline stage. **D.** Same as C for gamma coherence.

*p<0.05, **p<0.01, ***p<0.001, ****p < 0.0001, post-hoc Sidak's multiple comparisons following detection of main effects by Two-way ANOVA.

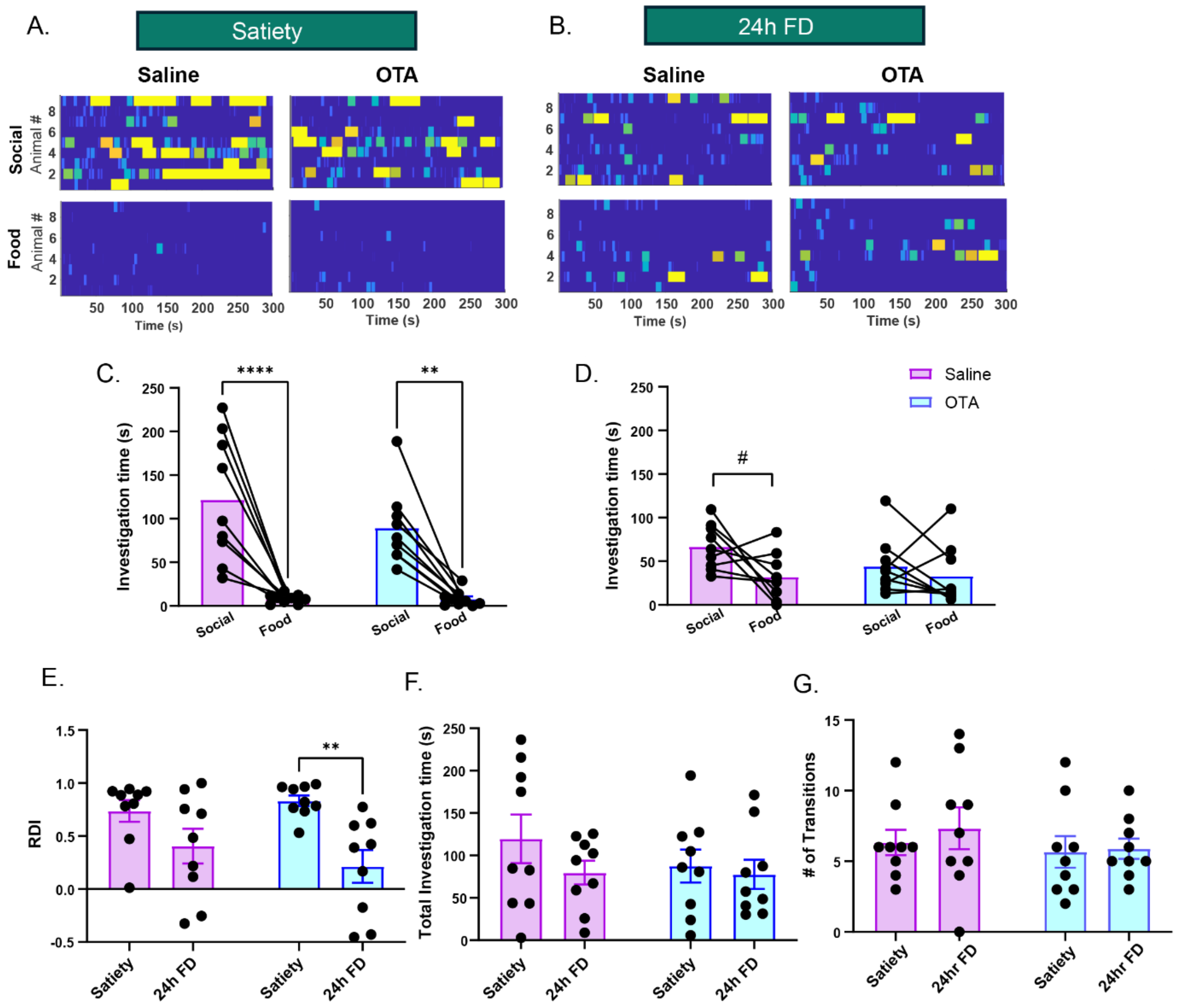


**Fig S3. Behavioral differences between surgically implanted rats in the saline and OTA groups in social vs food task. A-B.** Heatmaps of investigation bouts towards social or food stimulus between saline and oxytocin receptor antagonist (OTA)-treated rats at satiety in A, 24 hours of food-deprivation (24h FD) in B **C.** Statistical comparison between the social and food investigation time between saline and OTA-treated rats at satiety. **D.** Same as C for 24h FD. **E-G.** relative differential index (RDI) in G, total investigation time in H, number of transitions in I between saline and OTA-treated rats.

#p = 0.056, *p<0.05, **p<0.01, ****p<0.0001, post-hoc Sidak's multiple comparisons following detection of main effects by Two-way ANOVA.

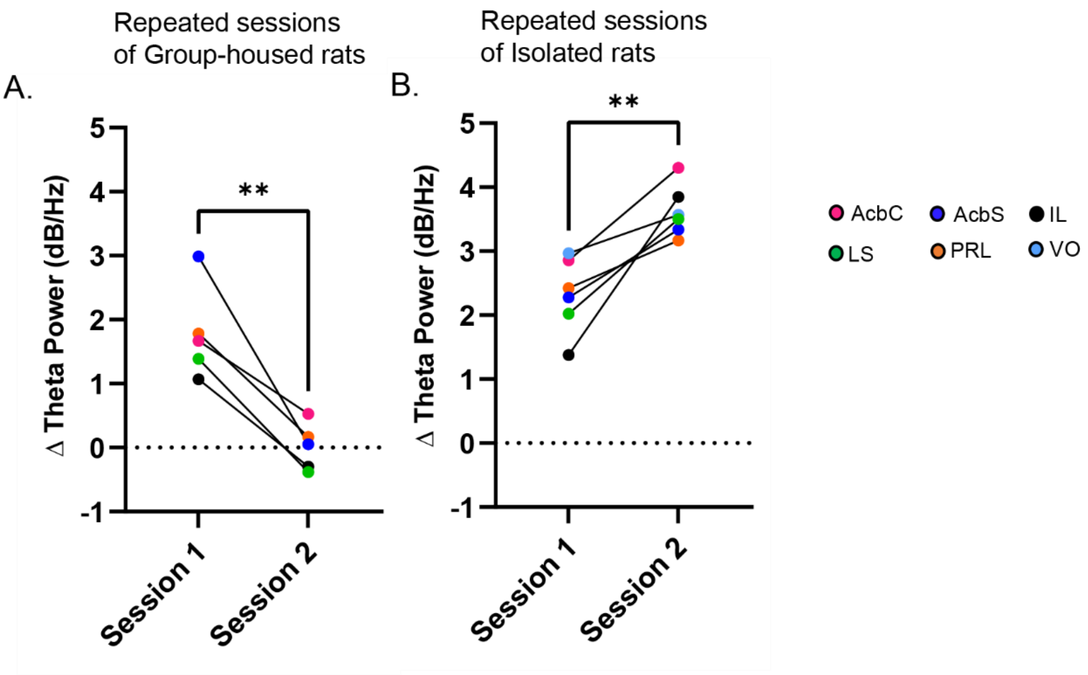


**Fig S4. Theta power differences across repeated sessions of social vs food task. A.** Comparison of change in theta power between the first and second session of SVF task recorded at satiety from the same grouped rats. **B.** Comparison of change in theta power between the first and second session of SVF task recorded at satiety from the rats that were treated with saline after 24h FD for the first sessions of recordings.

**p<0.01 Unpaired t-test.
